# Bioactivity foundation model for hit identification and hit-to-lead optimization from sequence data

**DOI:** 10.64898/2026.10.26.762100

**Authors:** Huan Yee Koh, Harvey Jia Wei Koh, Yizhen Zheng, Vivek Naranbhai, Daniel F. Schmidt, Johan Barthelemy, Ying-Ja Chen, Anh T.N. Nguyen, Shirui Pan, Lauren T. May, Geoffrey I. Webb

## Abstract

Learning from large-scale bioactivity archives is complicated by heterogeneous assay readouts and differences in data reliability. Here we show that a model that predicts probability distributions over binding affinities from sequence data opens model training to vast and diverse troves of heterogeneous bioactivity data. We develop PSICHIC+, a bioactivity foundation model trained on approximately 82 million protein–ligand records using a distributional objective that preserves quantitative resolution while incorporating variation in data reliability. Given a protein sequence and ligand SMILES, PSICHIC+ predicts binding affinity in milliseconds. In zero-shot virtual-screening benchmarks without target-specific training data, it improves enrichment over regression- and classification-based approaches and performs competitively with leading structure-based methods. In lab-in-the-loop campaigns, distribution-guided acquisition improves recovery of high-activity compounds under fixed screening budgets. Combining PSICHIC+ with Boltz-2 improves hit-to-lead affinity prediction over Boltz-2 alone, with mean correlation approaching that of FEP+ on the evaluated benchmarks, without requiring high-resolution protein structures. These results support distributional learning as a scalable approach to using heterogeneous bioactivity data for hit identification and hit-to-lead optimization.

## Introduction

Identifying small-molecule ligands with measurable activity against target proteins is central to chemical biology and drug discovery.^1,2^ Computational prediction of protein–ligand interactions supports both hit identification, which requires rapid screening of large chemical libraries to identify promising starting points, and hit-to-lead optimization, which requires accurate prioritization of newly designed candidates within and across chemical series. Current approaches occupy different operating regimes: structure-based and physics-based methods can achieve high accuracy but depend on explicit structural modeling and substantial computation, whereas sequence-based predictors avoid structural prerequisites and scale readily but have not fully exploited large, heterogeneous bioactivity repositories indexed by protein and ligand sequence. Here we show that representing uncertainty as a conditional distribution over bioactivity measures allows noisy, heterogeneous sequence data to be used directly, rather than discarded or reduced to binary labels, while preserving the quantitative resolution needed to rank close analogs. We introduce PSICHIC+, a sequence-based bioactivity foundation model trained on approximately 82 million bioactivity training records with beta distributional outcomes. Sequence-based PSICHIC+ performs affinity prediction at millisecond scale, approximately 15,000-fold faster than Boltz-2^3^, while an uncertainty-weighted PSICHIC+–Boltz-2 ensemble approaches FEP+-level^4^ accuracy at approximately 4,000-fold lower per-pair runtime (Fig. 1).

**Fig 1.**
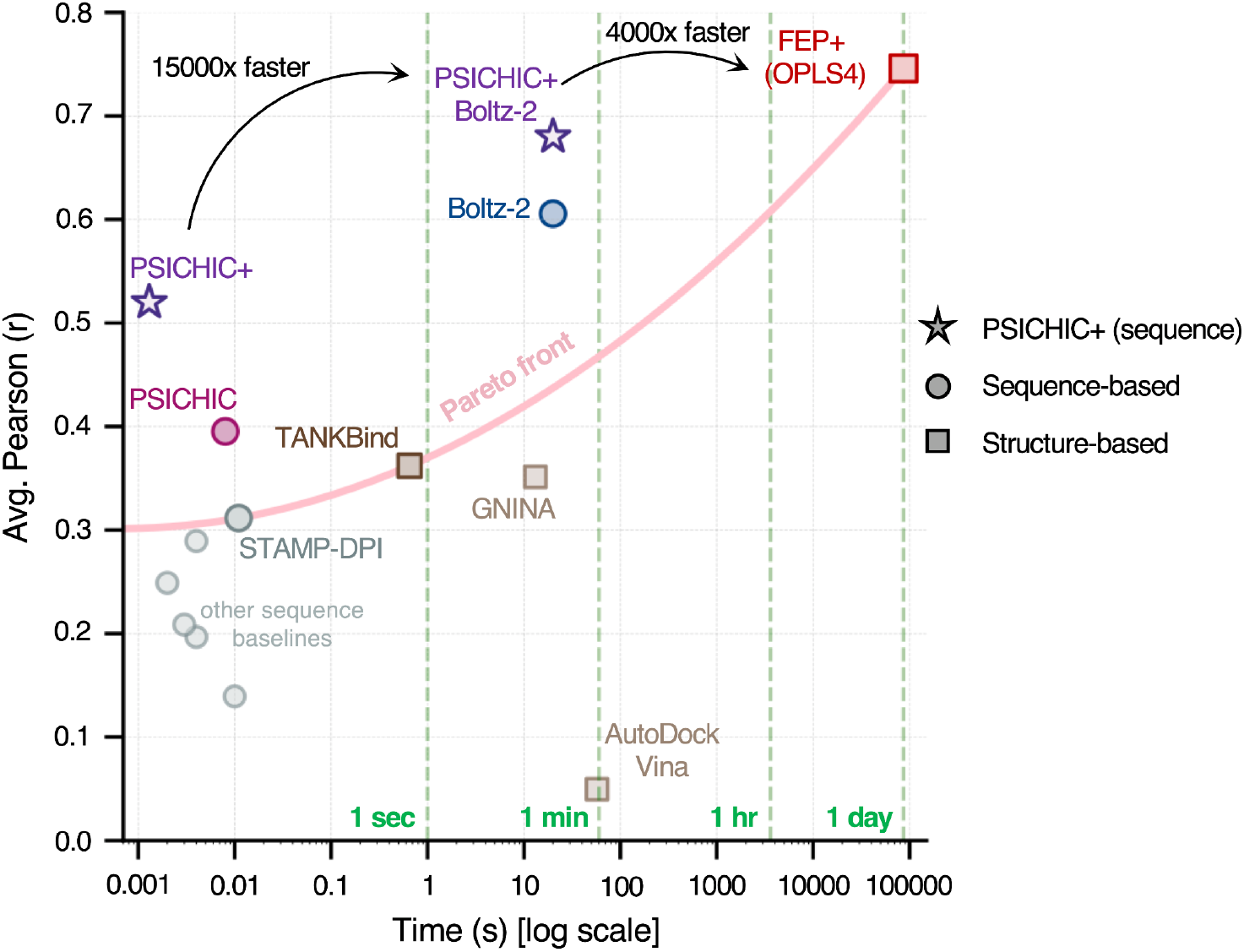
PSICHIC+ accuracy-speed trade-off for binding affinity prediction. Mean Pearson correlation (r) is plotted against wall-clock inference time per protein–ligand pair on a log scale. Stars denote PSICHIC+ variants, circles denote other sequence-based baselines and squares denote structure-based methods. All sequence-based models were evaluated on the same NVIDIA A100 GPU hardware. AutoDock Vina, a structure-based docking method, was run on CPU. PSICHIC+ reduces per-pair inference time to 1.34 ms, a 5.98-fold improvement over the original PSICHIC implementation, through NVIDIA GPU optimizations that include BioNeMo-based input reuse^5^ (see Methods and Supplementary Fig. 1). The pink curve represents a hypothetical Pareto front fitted to STAMP-DPI, TANKBind and FEP+ (OPLS4); vertical dashed lines indicate 1 s, 1 min, 1 h and 1 day. Runtime annotations indicate the reference method for each comparison: sequence-only PSICHIC+ is approximately 15,000× faster than Boltz-2^3^ based on runtimes of 1.34 ms and approximately 20 s per pair, whereas the PSICHIC+–Boltz-2 ensemble is approximately 4,000× faster than FEP+ based on runtimes of approximately 20 s and approximately one day per pair.^3^

Experimental screening and DNA-encoded libraries^6^ (DELs) can interrogate large molecular collections experimentally, but such campaigns are costly, assay-dependent and difficult to repeat across many targets. Computational prioritization can reduce this experimental burden, and many established workflows are structure-guided,^7–11^ using experimentally determined or predicted protein structures for docking and scoring, structural interpretation of structure–activity relationships and free-energy calculations.

The strength of structure-based approaches lies in their physical and geometric grounding, but the same dependence creates their principal bottlenecks. Performance relies on accurate structures, relevant protein conformations, reliable binding-site definitions and tractable ligand-pose sampling.^7^ Insilico prediction of structures has expanded structural coverage,^12^ but does not fully resolve protein dynamics, induced fit or assay-relevant conformational states.^13–15^ Co-folding models such as Boltz-2 broaden this paradigm by predicting a bound protein–ligand complex from sequence inputs and coupling the inferred geometry to affinity prediction. Yet they remain structure-first: a detailed three-dimensional complex must be generated for every protein–ligand pair before it can be scored. This adds substantial per-pair computation and allows errors in protein conformation, binding-site localization or ligand pose to propagate into the affinity estimate. More accurate physics-based methods impose still greater computational cost, limiting exhaustive screening of very large libraries and rapid design–make–test cycles.

This limitation matters because most experimental data in ligand discovery is not structural. High-quality structural repositories, including the Protein Data Bank, remain at the 10^5^–10^6^ scale,^11,16–18^ whereas public bioactivity repositories contain 10^7^ to 10^8^ protein–ligand measurements,^7,19,20^ with proprietary collections substantially larger still. These records pair protein sequences and ligand molecular strings with binding, potency and screening readouts, usually without a resolved binding site, ligand pose or bound complex. A structure-first pipeline must therefore reconstruct geometry that the underlying experiment did not measure. At repository scale, retaining uncertain reconstructions propagates structural errors, whereas filtering them can disproportionately remove sparsely studied targets and uncommon chemotypes. Sequence-based models offer a different operating regime: they learn directly from protein sequences, ligand molecular strings and assay readouts, allowing the much larger bioactivity corpus to serve as supervision without first converting every record into a predicted complex.^21^ In this setting, protein–ligand modeling suggests a data-scaling corollary of Sutton’s bitter lesson^22^: methods able to learn directly from the largest task-relevant data source may capture predictive signal unavailable to approaches constrained to smaller, structurally resolved corpora.

Scale alone, however, is not enough. Public bioactivity repositories combine direct affinity measurements, functional potency readouts, high-throughput screening labels, assay-dependent activity calls and synthetic or inferred negatives, with substantial variation in units, protocols and reliability.^20,23^ Existing sequence-based learning strategies typically resolve this heterogeneity in one of two ways. Regression models apply stringent curation to retain comparable quantitative measurements, preserving activity resolution but discarding much of the available corpus.^7,23^ Classification models retain broader coverage by thresholding diverse readouts into active or inactive labels, but erase the quantitative distinctions needed to rank compounds within and across chemical series.^11,19^ The central challenge is therefore to exploit the scale of heterogeneous bioactivity data without sacrificing the quantitative resolution required for hit-to-lead optimization.

One response has been to treat individual assays as separate learning tasks and transfer knowledge across them, enabling rapid adaptation to a new assay from a small set of assay-specific measurements.^24^ Although such models can resolve subtle activity differences within a chemical series, deployment in a new discovery program still requires target- and assay-specific measurements. In effect, assay-specific adaptation forgoes the goal of a broadly deployable bioactivity foundation model. This shift raises an important question: can a sequence-based bioactivity foundation model learn directly from large-scale, noisy and assay-inconsistent data while retaining the resolution needed to decode subtle activity differences in protein–ligand interactions? PSICHIC+ brings this sequence-first scaling argument to its ultimate conclusion: a single bioactivity foundation model trained directly on heterogeneous measurements can support library-scale, zero-shot hit identification while retaining the quantitative resolution needed to rank candidate compounds within and across chemical series during hit-to-lead optimization.

## Results

### Development of a bioactivity foundation model

To learn from heterogeneous sequence-based bioactivity data without discarding quantitative measurements or reducing all readouts to binary labels, we developed PSICHIC+, a sequence-based bioactivity foundation model for protein–ligand interaction prediction (Fig. 2a). PSICHIC+ extends the existing PSICHIC architecture,^7^ which encodes physicochemical protein–ligand interaction patterns from protein sequence and ligand SMILES.

**Fig 2.**
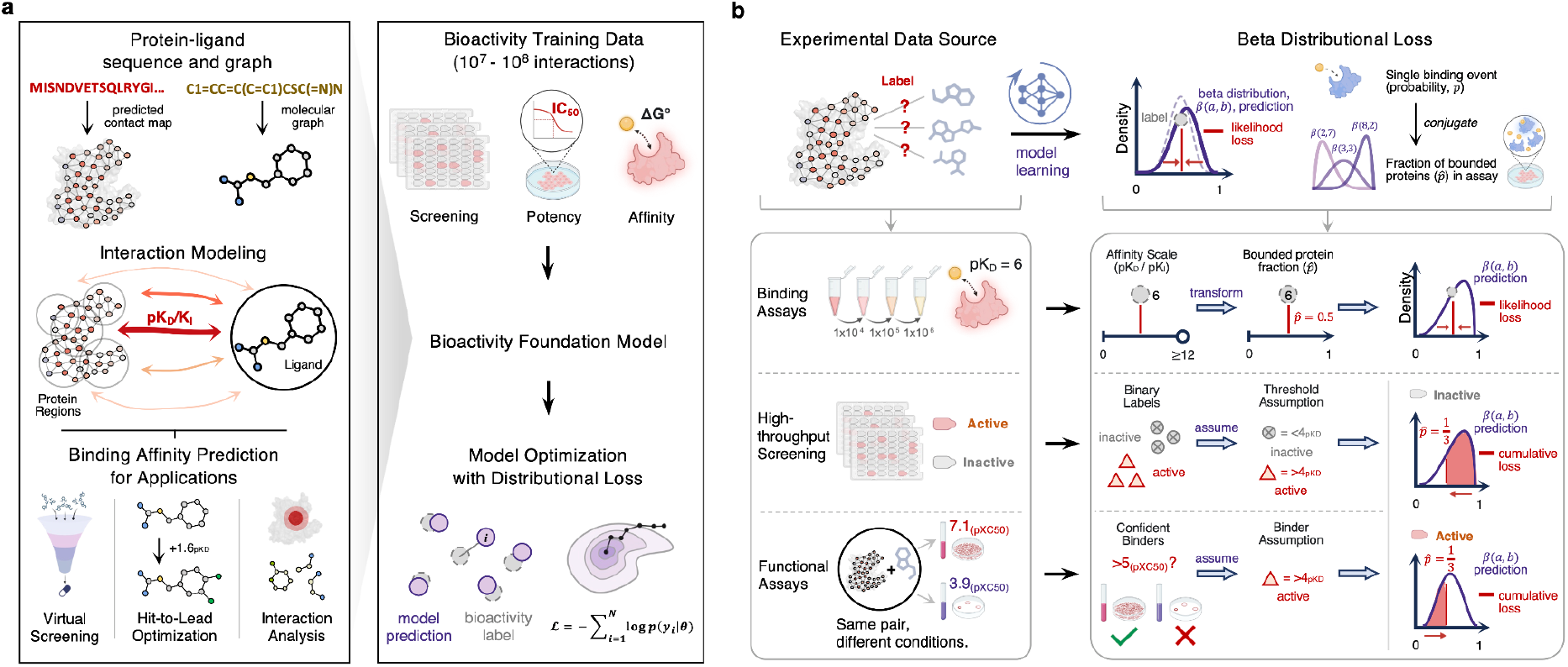
A bioactivity foundation model for hit identification and hit-to-lead optimization from sequence data. **a**, Schematic of PSICHIC+. A bioactivity foundation model for protein–ligand interaction prediction is trained on large-scale, sequence-anchored bioactivity corpora comprising approximately 82 million protein–ligand training records. Given a protein sequence and ligand SMILES, PSICHIC+ learns physicochemical protein–ligand interaction fingerprints and predicts a beta distribution over normalized bioactivity, providing both an expected activity and a concentration parameter for the predictive distribution. These outputs support virtual screening, hit-to-lead optimization and interaction analysis through learned protein–ligand interaction fingerprints (Supplementary Figs. 2 and 3). **b**, Heterogeneous supervision from direct binding measurements, functional potency measurements and HTS-derived activity labels, including both binding- and function-based screens, provides broad coverage across protein targets, ligand chemotypes and assay readout types. The source-aware beta-distributional loss maps continuous and binary supervision into a shared probabilistic space and regularizes concentration according to assay/source quality and reliability, improving robustness to noisy labels and dataset shifts.

However, PSICHIC produces point predictions and is trained with either least squares regression or classification objectives. This separates training into two restricted regimes: curated continuous measurements for least squares regression, or active/inactive labels for classification. Least squares regression preserves affinity magnitude but cannot naturally incorporate HTS-derived labels or thresholded potency outcomes; it also outputs only a single point estimate that represents the mean of the expectation with no indication of expected precision. Classification can use broader supervision only after converting quantitative measurements into binary labels, removing potency magnitude and assay-specific variation.

PSICHIC+ removes this restriction by replacing deterministic regression or classification heads with a beta-distribution output head. For each protein sequence and ligand SMILES pair, PSICHIC+ predicts the two beta shape parameters α and β over normalized bioactivity (Fig. 2b), where both parameters, α and β, define the beta distribution *Beta*(α, β). Specifically, each assay readout is treated as evidence about the same bounded bioactivity response. Exact pK-scale measurements are transformed to the unit interval and trained with a likelihood loss on the observed value. Binary or thresholded evidence, including HTS active/inactive calls, functional potency labels and generated negatives, supervises the probability that activity exceeds a predefined threshold through the beta cumulative distribution function. In this manner, PSICHIC+ can report the beta mean, 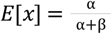, rescaled to the pK range, as a point prediction. The concentration κ = α + β controls the width of the predictive distribution. For a fixed mean, the variance is *Var*(*X*) = *E*[*X*](1 − *E*[*X*])/(κ + 1), and a high concentration means that the probability mass is tightly concentrated around the point estimate, suggesting that a realized assay value is more likely to be close to that estimate. This allows coarse labels to inform the activity landscape without forcing such labels into artificial point estimates.

We trained PSICHIC+ on a large-scale corpus curated from Papyrus,^19^ integrating direct affinity measurements, functional potency readouts and high-throughput screening labels (see Methods) across diverse proteins, organisms and chemotypes (see Supplementary Fig. 4). We also generated per-target decoy molecules as synthetic negative examples (see Methods). The final training corpus comprised approximately 82 million records spanning 4,900 unique proteins and approximately 6.1 million unique ligands.

**Fig 3.**
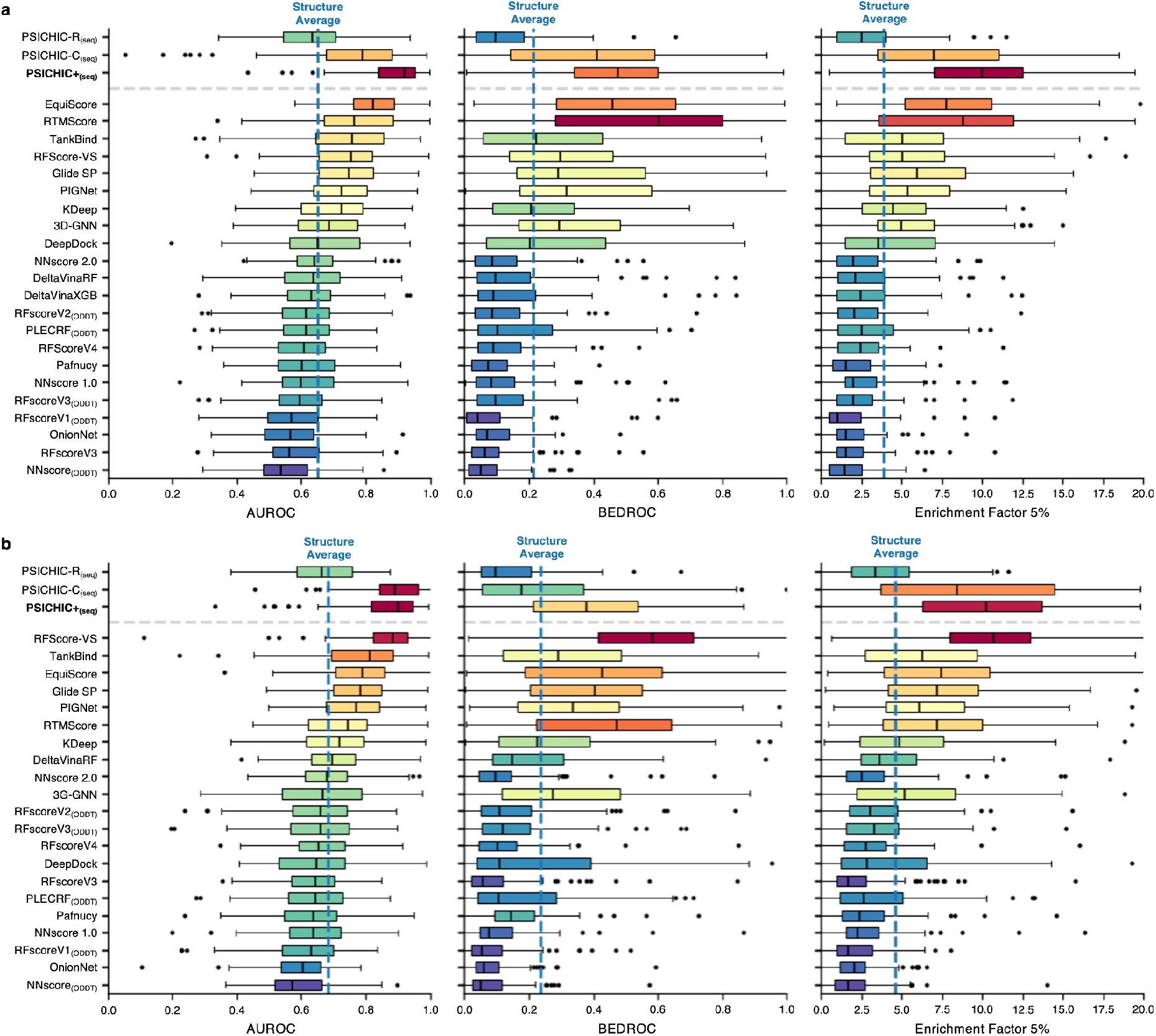
Benchmarking PSICHIC+ for virtual screening on DUD-E and DEKOIS 2.0. **a**,**b**, Boxplots show target-level virtual-screening performance on DUD-E (**a**, 102 targets) and DEKOIS 2.0 (**b**, 81 targets) for AUROC, BEDROC (α = 80.5) and enrichment factor at the top 5% of ranked compounds (EF5%). The center line denotes the median across targets, box limits indicate the interquartile range, whiskers extend to 1.5× the interquartile range and points denote outlier targets. Box colors encode median performance ranging from blue for lower relative performance to red for higher relative performance. The top three rows are sequence-based PSICHIC variants using only protein sequence and ligand SMILES: least-squared-regression-trained PSICHIC-R, classification-trained PSICHIC-C and distributional PSICHIC+. All methods below the gray dashed line are structure-based comparators that use protein structural information. Boltz-2 was not included in this comparison because retraining the co-folding model under the same 90% sequence-identity exclusion protocol would be computationally prohibitive. The blue dashed line marks the average performance of the structure-based methods. PSICHIC+ improves early-retrieval performance over both PSICHIC baselines and performs competitively with leading structure-based virtual-screening methods.

**Fig 4.**
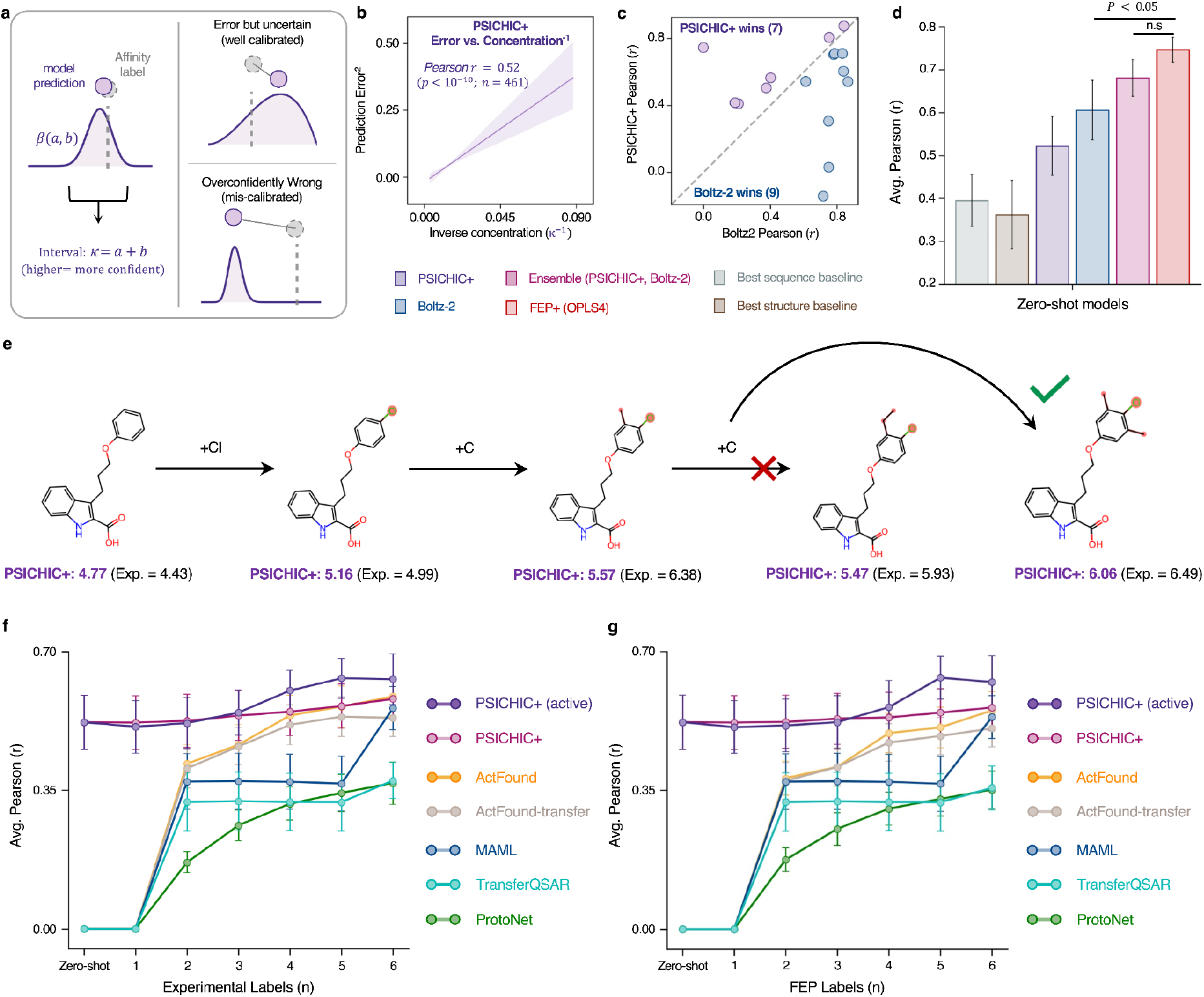
Concentration-aware affinity prediction and label-efficient adaptation with PSICHIC+. **a**, PSICHIC+ predicts a beta distribution β(α, β) over affinity and reports concentration through the concentration κ = α + β: higher κ denotes narrower central predictive intervals. Schematic examples show narrow distributions with small errors (top right) and overly narrow distributions with large errors (bottom right). **b**, Across held-out protein–ligand pairs, prediction error increases with the inverse of concentration (Pearson *r* = 0. 52, *P* < 10^−10^; *n* = 461), showing that concentration tracked empirical prediction error. The shaded band represents the 95% CI of the fit. **c**, Head-to-head zero-shot accuracy of PSICHIC+ versus a strong structure-based baseline (Boltz-2) by protein; points above the diagonal favor PSICHIC+. **d**, Bars, from left to right, denote the best sequence-only baseline (PSICHIC)^7^, the best structure-based deep learning baseline (TANKBind)^25^, PSICHIC+, Boltz-2^3^, the concentration-modulated PSICHIC+–Boltz-2 ensemble and FEP+ (OPLS4)^4^. Error bars denote s.e.m. across targets. Statistical comparisons used one-sided Wilcoxon signed-rank tests on per-target correlations. The two reported P values were not adjusted for multiple comparisons. The PSICHIC+–Boltz-2 ensemble is the only model not significantly outperformed by FEP+. FEP+ exceeds all non-ensemble baselines (*P* < 0. 05; one-sided Wilcoxon signed-rank test on per-target correlations; *n* = 16). **e**, Example of PSICHIC+ zero-shot prediction of chemistry trajectory: small, chemically plausible edits (e.g., +Cl, carbon extension) progressively improve predicted affinity and match experimental measurements (purple: PSICHIC+; gray: experiment). **f**,**g**, Few-shot adaptation with experimental labels (**f**) or FEP+ labels (**g**). PSICHIC+ starts from the strongest zero-shot baseline and remains highest across small support sets. Additional labels provide smaller marginal gains because the pretrained model is already informative. Active selection further improves PSICHIC+ over random support selection, indicating concentration-guided refinement of an already strong prior. Curves show mean Pearson correlation across proteins with s.e.m. error bars.

Rather than treating every label as equally informative, PSICHIC+ makes source-specific label quality and reliability part of the learning objective. In the implemented concentration parametrization, the prediction defines the interior mode of the conditional beta distribution, while the fitted beta mean is computed from the resulting shape parameters. PSICHIC+ therefore regularizes concentration according to data source: higher-quality affinity measurements are assigned a higher reference concentration, whereas HTS labels, synthetic negatives and other weakly supervised sources are assigned lower reference concentrations (see Methods). This discourages noisy or coarse labels from driving the model towards implausibly narrow predictive distributions, while still allowing them to shape the learned activity landscape. The resulting objective is a quality-aware distributional learning framework that integrates binding measurements, functional potency readouts, screening-derived labels and decoys in a shared probabilistic space, with each prediction accompanied by a predictive distribution whose concentration is regularized based on assay quality and variability across heterogeneous data sources.

### Hit Identification with PSICHIC+

We evaluated PSICHIC+ for virtual screening against the standard PSICHIC model and benchmarked both against 22 structure-based methods on DUD-E^8^ and DEKOIS 2.0^9^. To assess generalization to unseen targets, we retrained all PSICHIC variants separately for each benchmark after removing from the curated training corpus any proteins sharing at least 90% sequence identity with a benchmark target.

PSICHIC+, trained with a distributional loss, consistently outperformed variants trained using either least-squared point regression or classification (Fig. 3). The largest gains were observed in early-enrichment metrics, which are particularly important in virtual screening. On DUD-E, PSICHIC+ achieved an EF5% of 13.3 and a BEDROC of 0.51, compared with 5.6 and 0.10 for the point-regression model and 10.8 and 0.34 for the classification model, respectively. EF5% measures the enrichment of active compounds within the top 5% of the ranked library, whereas BEDROC places greater emphasis on actives retrieved near the top of the ranking. PSICHIC+ showed the same advantage on DEKOIS 2.0, outperforming the point-regression and classification variants across all evaluation metrics. These findings suggest that representing bioactivity as a predictive distribution improves the model’s ability to distinguish among protein–ligand pairs and prioritize active compounds early in the ranked list.

PSICHIC+ also compared favorably with the 22 structure-based virtual-screening methods. On DUD-E, it outperformed every structure-based comparator across AUROC, BEDROC and EF5%. On DEKOIS 2.0, its performance approached that of the two strongest structure-based methods, RTMscore^10^ and Equiscore^11^, with EF5% values of 8.0, 8.1 and 8.2, respectively. This performance is notable because PSICHIC+ requires only a protein sequence and ligand SMILES, whereas structure-based methods depend on experimentally determined three-dimensional protein structures and more computationally intensive docking or scoring workflows.

Together, these results show that the distributional objective improves virtual-screening performance over the PSICHIC point-prediction variants. By matching and even exceeding established structure-based pipelines while retaining rapid, structure-free inference, PSICHIC+ is well suited to large-scale screening, particularly for targets lacking high-quality structural information.

### Hit-to-lead optimization with PSICHIC+

Hit-to-lead optimization refines promising ‘hit’ chemotypes into lead compounds with improved affinity, potency, selectivity, and other pharmacological properties. A key challenge in this process is series ranking: distinguishing subtle structural variations among closely related analogs that substantially influence pharmacological properties, in order to prioritize synthesis and candidate progression. This phase has traditionally been driven by physics-based free-energy simulations such as free energy perturbation (FEP)^26^, together with docking and structure-guided structure–activity relationship (SAR) analysis.^7^ Assay-labeled data can also support chemistry-driven SAR analysis and target-specific machine-learning models^24^. However, these data are rarely available at project outset because such models require labeled analogs measured for the same target under the same assay conditions. As a result, early design–make–test cycles often lag or cannot proceed at the desired pace while high-resolution structures and chemistry-based datasets are still under development^27^.

PSICHIC+ overcomes these structural and labeling bottlenecks by learning directly from heterogeneous bioactivity corpora and transferring interaction patterns across targets. After pretraining, PSICHIC+ outputs a beta distribution, Beta(α, β), over affinity and reports its concentration parameter κ = α + β. A larger κ implies narrower predictive intervals. The model therefore provides an affinity estimate and a concentration parameter for each protein–ligand pair without requiring structural inputs or target-specific labels (Fig. 4a). We evaluated PSICHIC+ on established FEP benchmark sets (JACS^28^ and Merck^29^; 16 unique target assays with 29 compounds on average) in a zero-shot setting after excluding all benchmark ligands and proteins from pretraining (see Methods). Prediction error increased as concentration parameter of PSICHIC+’s beta predictive distribution decreased (Pearson *r* = 0. 52, *P* < 10^−10^; *n* = 461), showing that high-concentration predictions were more likely to lie close to the realized activity value, whereas low-concentration predictions flagged cases where the point estimate should be treated with caution. This makes the predictive distribution useful for triaging designs, prioritizing costly synthesis and deciding when additional validation is needed (Fig. 4b).

To assess accuracy, we benchmarked PSICHIC+ against zero-shot sequence-based baselines, structure-based baselines, Boltz-2^3^ (an AlphaFold 3-class^21^ affinity predictor), and FEP+ (OPLS4)^4^. PSICHIC+ surpassed Boltz-2 on 7 of 16 targets (Fig. 4c) and outperformed the leading sequence- and structure-based baselines, while FEP+ had the highest mean Pearson correlation, at 0.74 (Fig. 4d). Although Boltz-2 attained a higher overall correlation than PSICHIC+ (0.60 vs. 0.52), performance varied across targets. PSICHIC+ concentration correlated with its own error, so we used it to modulate a simple ensemble with Boltz-2. The ensemble starts from equal weights and reduces the PSICHIC+ contribution only when the models agree and PSICHIC+ concentration is low (see Methods). This PSICHIC+–Boltz-2 ensemble raised the mean zero-shot correlation to 0.68 and was the only method not significantly outperformed by FEP+ (Fig. 4d). These findings establish PSICHIC+ as a robust, structure-free complement to physics-based workflows, particularly when structures are unavailable or high-throughput prioritization is essential.

We further examined PSICHIC+ performance in a few-shot adaptation setting by fine-tuning an analog-ranking model using a modest target-specific support set consisting of a few labeled analogs from the same target and assay. For each target, we varied support size from *N* = 1 to *N* = 6 and tested two PSICHIC+ variants: PSICHIC+ (random), which samples support compounds uniformly at random, and PSICHIC+ (active), which selects compounds using an upper-confidence-bound acquisition score that combines the predictive mean and beta standard deviation (see Methods). We compared these against five meta-learning or transfer-learning baselines designed to learn from limited assay-labeled data: MAML^30^, ProtoNet^31^, TransferQSAR, ActFound-transfer^24^ and ActFound^24^. Unlike PSICHIC+, these baselines lack zero-shot capability, requiring a support set (*N* ≥ 2) before inference. Across all *N*, PSICHIC+ (random) outperformed every baseline, with ActFound nearing equivalence only at higher *N*; PSICHIC+ (active) delivered further gains at equivalent label budgets (Fig. 4f). This trend persisted with FEP-derived labels in place of experimental ones (Fig. 4g), where PSICHIC+ (active) with FEP-derived labels matched the performance it achieved with experimentally-derived labels. These results support a hybrid workflow in which a limited set of FEP computations can seed the optimization process, enabling PSICHIC+ to be fine-tuned on limited data and rank any number of analogs at very low additional inference cost.

Taken together, PSICHIC+ provides structure-free, zero-shot prioritization when no target-specific data exist, and label-efficient few-shot adaptation when only a handful of analogs can be measured. The predictive distribution can also be used both to arbitrate predictions and to select compounds for labeling, focusing experimental or FEP measurements on candidates expected to provide the most information. A limited set of high-quality labels can then support rapid scoring of thousands to millions of analogs. Thus, in practice, only a modest number of high-quality labels is required to bootstrap PSICHIC+, after which inference scales to thousands of analogs at very low additional inference cost. This creates economies of scale that support rapid iteration and makes PSICHIC+ a complement to more computationally demanding models like Boltz-2 and to physics-based pipelines.

### Lab-in-the-Loop Optimization with PSICHIC+

To extend PSICHIC+ beyond one-off zero- or few-shot applications, we embed it in a lab-in-the-loop workflow^32^. Medicinal chemistry proceeds through iterative “design-make-test” cycles; therefore, the most effective models are those that continuously learn from feedback rather than relying on static snapshots. Lab-in-the-loop refers to an iterative process in which the model proposes compounds using an uncertainty-guided acquisition function. Once a tractable batch is assayed, the newly labeled data are used to fine-tune the model for the subsequent round of proposals. This closed-loop approach is valuable because it concentrates measurements where they are most informative, adapts to assay-specific nuances or evolving SAR, and systematically balances the exploitation of high-scoring candidates with exploration of under-sampled regions.

PSICHIC+’s lab-in-the-loop procedure is as follows. First, we score the unlabeled pool to obtain the predictive mean and concentration (Fig. 5a, step 0). We then rank candidates by the acquisition function and select a batch for labeling (Fig. 5a, step 1), either prioritizing the highest-affinity compounds (PSICHIC+ Greedy) or balancing exploitation with uncertainty-guided exploration to maximize information gain (PSICHIC+ upper-confidence-bound (UCB); see Methods). The selected batch is assayed to obtain labels (Fig. 5a, step 2), appended to the training set (Fig. 5a, step 3), and the model is fine-tuned on all accumulated labels; calibration is refreshed, and the next cycle begins. As labels accumulate, each cycle updates the ranking. In this setting, the objective is to recover the highest-affinity compounds from a large virtual library under a fixed labeling budget, thereby accelerating the discovery of potent chemotypes. Because PSICHIC+ can score compounds before target-specific labels are available, the first cycle can begin without the random initial sample required by conventional target-specific models (Fig. 5a).

**Fig 5.**
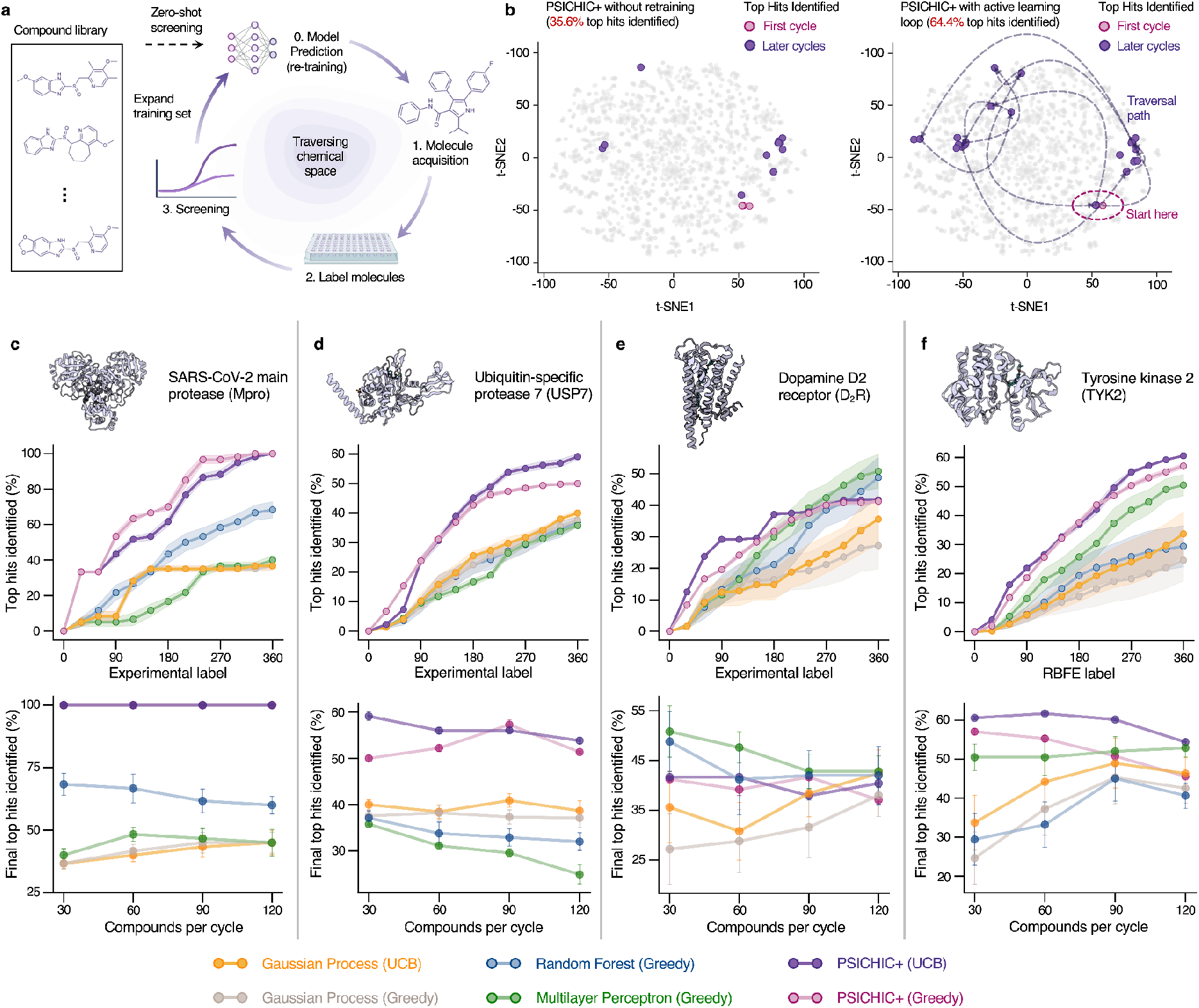
Lab-in-the-loop PSICHIC+ workflow accelerates hit-to-lead optimization. **a**, Iterative active-learning workflow in which PSICHIC+ performs zero-shot screening of a compound library, selects 30–120 candidates per cycle using greedy or upper-confidence-bound (UCB) acquisition, obtains labels, expands the training set and retrains before the next cycle. **b**, t-SNE embeddings of the USP7 compound library showing selected compounds across acquisition cycles for PSICHIC+ without retraining (left; 35.6% top hits identified) and PSICHIC+ with active-learning retraining (right; 64.4% top hits identified). The percentage of top hits identified is the proportion of the library’s true top-1% compounds selected within the fixed labeling budget. **c–f**, Active-learning performance for SARS-CoV-2 main protease (Mpro) (**c**), ubiquitin-specific protease 7 (USP7) (**d**), dopamine D2 receptor (D2R) (**e**) and tyrosine kinase 2 (TYK2) (**f**). Upper panels show cumulative top hits identified as labels acquired over successive 30 sample batch cycles; lower panels show final top hits identified after multiple cycles for batch sizes of 30, 60, 90 and 120 compounds per cycle until the total budget reaches 360 labels. Top hits are defined by experimental activity or affinity labels for Mpro, USP7 and D2R, and by RBFE labels for TYK2. PSICHIC+ with greedy or UCB acquisition identifies more top hits than random forest, multilayer perceptron and Gaussian process baselines across most targets and batch sizes.

To assess the benefit of lab-in-the-loop iteration over a static screen, we compared PSICHIC+ operating in a lab-in-the-loop setting with a single PSICHIC+ zero-shot pass for recovering the highest-affinity compounds (defined as the top 1% by affinity within each dataset). As an illustrative case from our benchmark suite, we fixed the budget at 360 queries in batches of 30 compounds per cycle. On the therapeutically relevant ubiquitin-specific protease 7 (USP7) dataset, the lab-in-the-loop procedure recovered 64.4% of the highest-affinity compounds, whereas a static zero-shot ranking recovered only 35.6% (Fig. 5b). We then generalized this analysis by simulating lab-in-the-loop campaigns on four therapeutically relevant targets using curated affinity datasets^33^: SARS-CoV-2 main protease (Mpro)^34^, USP7^35^, dopamine D_2_ receptor (D_2_R)^36^, and TYK2 kinase^37^. Each dataset comprises chemically diverse compounds annotated with affinity labels (Mpro: *N* = 665; USP7: *N* = 4, 535; D_2_R: *N* = 2, 502; TYK2: *N* = 9, 997). All targets have experimental affinity values, except TYK2, for which labels derive from relative binding free energy (RBFE) calculations. For each target, we iterated fixed batches of 30 candidates selected by the acquisition rule until 360 molecules had been evaluated, enabling controlled comparisons under the same labeling budget (Fig. 5c–f).

Across the four targets and tested batch sizes, PSICHIC+ recovered more top-affinity compounds than the baselines in most settings, ranking first for Mpro, USP7 and TYK2 (Fig. 5c–f, upper panels). Because laboratory throughput varies with assay format and plate capacity, we further evaluated batch sizes of 30, 60, 90, and 120 compounds per cycle until reaching the same total budget of 360 labels. PSICHIC+ maintained the most consistent performance across all batch sizes, indicating robustness to practical experimental constraints (Fig. 5c–f, lower panels). Across the four benchmark datasets, PSICHIC+ recovered more top-ranked compounds than the tested baselines in most target and budget settings. In contrast, complex predictors such as Boltz-2 use substantially more compute and are not designed for rapid assay-specific retraining; reported configurations use 128 A100 GPUs^3^, illustrating the difference in training scale. Structure-based methods also require protein structures and incur higher per-compound computational costs. PSICHIC+ fine-tuning used less than 10 GB of GPU memory, and its low per-pair inference time supports repeated analog ranking on a single GPU, including freely accessible platforms such as Google Colab.

### Accuracy-speed trade-off of PSICHIC+

The practical value of a predictor in hit identification and hit-to-lead optimization depends on both predictive accuracy and computational throughput. To quantify the accuracy–speed trade-off against representative sequence-based and structure-based methods, we compared mean Pearson correlation for affinity prediction with wall-clock inference time per protein–ligand pair from FEP benchmark sets (JACS^28^ and Merck^29^), using identical GPU hardware for sequence models and published runtimes for structure-based comparators (Fig. 1). Across these methods, higher accuracy was generally associated with longer runtimes. The optimized sequence-only model achieved a mean Pearson correlation of 0.52 at 1.34 ms per protein–ligand pair, compared with 0.60 for Boltz-2 at approximately 20 s per pair, representing an approximately 15,000-fold reduction in runtime. A concentration-modulated ensemble with Boltz-2 increased correlation to 0.68 at approximately 20 s per pair, narrowing the gap to FEP+ at 0.74 while reducing the reported per-pair runtime from approximately one day to approximately 20 s, a roughly 4,000-fold difference.

Across the benchmark, PSICHIC+ achieved a favorable operating point, attaining a mean correlation of *r* = 0. 52 with a mean inference time of 1.34 ms per protein–ligand pair. Although PSICHIC+ retains the PSICHIC backbone and adds only a distributional concentration output, NVIDIA GPU optimizations and BioNeMo-based input reuse^5^ reduced inference time relative to the original PSICHIC implementation (see Methods and Supplementary Fig. 1). Combining PSICHIC+ with Boltz-2 through concentration-modulated ensembling increased performance to r = 0.68 at approximately 20 s per pair. Thus, sequence-only PSICHIC+ is approximately 15,000× faster than Boltz-2, while the PSICHIC+–Boltz-2 ensemble is approximately 4,000× faster than FEP+ and approaches the accuracy of FEP+ (Fig. 1). In practical terms, PSICHIC+ is on the order of *60 million*-fold faster than FEP+ for only a modest loss in predictive correlation, while PSICHIC+–Boltz-2 is approximately 4,000-fold faster than FEP+ and approaches the accuracy of FEP+.

These results map directly onto typical discovery constraints. Sequence-based PSICHIC+ supports library-scale triaging and rapid hypothesis testing at very low additional inference cost. The concentration-modulated PSICHIC+–Boltz-2 ensemble then offers a high-accuracy mode that remains fast enough for routine prioritization and can be reserved for subsets where added precision is desirable. Together with the lab-in-the-loop workflow, this accuracy-throughput flexibility enables programs to allocate expensive calculations and assays where they yield the greatest benefit while maintaining rapid iteration across large chemical spaces.

## Discussion

Drug discovery can be accelerated by models that can learn from noisy real-world data, operate directly from sequence and screen large chemical libraries efficiently. PSICHIC+ addresses this need by learning from heterogeneous bioactivity measurements without reducing quantitative data to binary labels, thereby accessing the wealth of sequence-based bioactivity data denied to previous methods. Together, the predictive mean and concentration support principled ranking and concentration-aware decision-making across hit identification and hit-to-lead optimization. These results show that PSICHIC+ can complement physics-based workflows when structures are unavailable or high-throughput prioritization is required.

PSICHIC+ achieves high accuracy from sequence-only inputs with millisecond-scale inference. It unifies early discovery tasks in one framework, delivering low-cost, flexible hit identification via zero-shot prioritization of active chemotypes across diverse targets, and hit-to-lead optimization through very fast, accurate analog ranking. Concentration can guide arbitration and optional ensembling with structure-informed Boltz-2 predictors where added precision is valuable. The framework is also adaptable: it learns effectively from a few target-specific labels and integrates naturally with lab-in-the-loop optimization to focus measurements where they are most informative. Because inference is fast and fine-tuning fits within modest GPU memory, PSICHIC+ supports billion-compound virtual-screening campaigns and broad hit-to-lead programs. Its coverage across protein families may also support selectivity profiling and off-target risk assessment at a scale that is impractical for more computationally intensive models or physics-based pipelines.

Several opportunities exist to further extend the capabilities of PSICHIC+. A key priority is to broaden the diversity of the training corpus. Incorporating high-throughput data from DNA-encoded library selections^6^, together with data from fragment-based^38^ and covalent screening campaigns^39^, could improve generalizability across target classes and binding modalities. In parallel, enriching assay metadata with explicit representations of cofactors, pH, isoform identity and experimental protocols would enable more precise modeling and concentration estimates. An important next step for receptor systems is to separate the pharmacological processes currently represented by a single bioactivity response. Functional potency reflects ligand affinity and efficacy, together with system-dependent signal amplification, including receptor reserve. A hierarchical extension of PSICHIC+, informed by the Black–Leff operational model of agonism and subsequent transduction frameworks developed by Kenakin and Christopoulos, could represent affinity, efficacy and system-dependent transduction as distinct but coupled latent factors.^40–42^ Estimating these components would require matched binding and full concentration-response data, together with metadata on receptor abundance, signaling pathway and cellular context. Such a model could separate affinity-driven potency from efficacy- and system-dependent responses and support pathway-aware hit-to-lead optimization. Distributional learning also provides a natural interface with generative chemistry. Coupling PSICHIC+ to a molecular generator could rank proposed candidates by expected potency, predicted concentration and the probability of exceeding a desired activity threshold. This approach could guide exploration toward uncertain but potentially high-potency chemotypes. When paired with rapid synthesis platforms, such a closed-loop framework could accelerate discovery across underexplored regions of chemical space.

## Methods

### Distributional formulation of normalized bioactivity

PSICHIC+ models heterogeneous protein–ligand measurements using a bounded bioactivity variable. For direct binding measurements, this formulation can be motivated by a simple equilibrium occupancy model. Under a 1:1 binding model at fixed ligand concentration, the state of an individual receptor is binary: it is either ligand-bound or unbound. The occupancy of a randomly sampled receptor can therefore be represented as a Bernoulli variable with success probability θ, where θ denotes the fraction of receptors bound. At equilibrium,

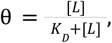

where [*L*] is ligand concentration and *K*_*D*_ is the dissociation constant. Thus, for a fixed reference concentration, lower *K*_*D*_, or equivalently higher *pK* _*D*_, corresponds to higher expected receptor occupancy. This occupancy interpretation provides a conceptual bridge between pK-scale binding measurements and a bounded, probability-like response variable.

In practice, PSICHIC+ does not assume that observed assay endpoints directly reflect equilibrium receptor occupancy. Large bioactivity corpora contain heterogeneous endpoints, including direct affinity measurements, functional potency readouts, HTS-derived activity labels and synthetic negative examples. We therefore use the equilibrium argument only as motivation for representing quantitative pK-scale measurements on a bounded interval. Direct pK-scale values are transformed to

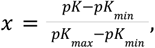

where *pK*_*min*_= 0 and *pK*_*max*_ = 12. This affine transformation maps pK-scale measurements to *x ∈* [0, 1], preserves their ordering and allows fitted beta means and central predictive intervals to be transformed back to pK units. Values are clipped away from the exact boundaries during training for numerical stability.

To model the conditional spread of this normalized response, PSICHIC+ predicts a beta distribution,

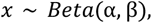

for each protein–ligand pair, with shape parameters conditioned on the protein and ligand inputs. The beta distribution is well suited to this setting because it is defined on the unit interval, can represent both broad and sharply concentrated fitted conditional response distributions, and supports interpretable location-and-concentration parameterizations. Specifically, the model predicts an interior beta location μ *∈* (0, 1) and a positive excess-concentration parameter φ > 0, which are converted to shape parameters as

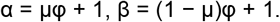

The excess-concentration parameter φ is the positive output of the prediction head. Because each shape parameter receives a unit offset, the total beta concentration is κ = α + β = φ + 2. The unit offset ensures α, β > 1 and therefore restricts the model to beta densities with a single interior mode. Numerical stability is handled separately by clipping boundary targets and using stable density and cumulative-distribution calculations. The reported point prediction is the fitted beta mean,

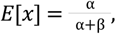

whereas the concentration α + β controls dispersion: a high concentration in the conditional distribution suggests that the point estimate is likely to be close to a realized assay value from that distribution. PSICHIC+ uses different likelihood terms depending on the type of supervision. For direct quantitative pK labels, the model minimizes the negative log-likelihood of the transformed value under the predicted beta density:

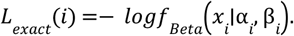

For labels available only as active/inactive outcomes relative to an activity threshold *T*, the threshold is first rescaled to

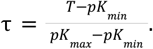

The model then predicts the probability that the latent response exceeds this threshold:

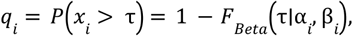

where *F Beta* is the beta cumulative distribution function. Binary labels are trained with cross-entropy on this threshold-exceedance probability:

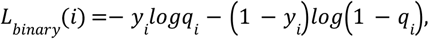

where *y*_*i*_ = 1 denotes an active label and *y*_*i*_ = 0 denotes an inactive label.

Equivalently, the supervision loss can be written as

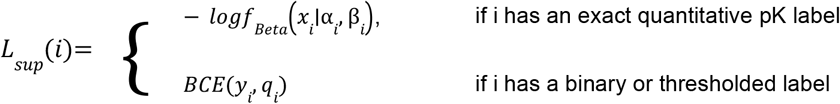

where *q* = *P* (*x*_*i*_ > τ = 1) − *F*_*Beta*_ (τ*∣*α _*i*_, β_*i*_) is the probability that the latent bioactivity response exceeds the threshold 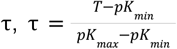 and *BCE*(*y*_*i*_, *q*_*i*_ ) =− *y*_*i*_ *logq*_*i*_ − (1 − *y*_*i*_ )*log*(1 − *q*_*i*_ ). Here, *f*_*Beta*_ and *F*_*Beta*_ denote the beta probability density function and cumulative distribution function, respectively. This formulation allows quantitative affinity measurements and binary or thresholded bioactivity labels to contribute to a shared distributional objective.

Finally, PSICHIC+ regularizes beta concentration according to data-source reliability. Each training example is assigned a source-quality tier: direct affinity measurements provide the most reliable supervision, whereas noisier sources, such as HTS-derived labels and synthetic negatives, receive lower reliability tiers. For each source tier *t*, we defined κ_*max*_ (*t*) from a prespecified reference 95% predictive-interval half-width *e*_*t*_ on the pK scale. We converted this value to the normalized scale as 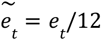 and used a symmetric beta reference distribution centered at 0.5. Specifically, the tier-specific reference was obtained by one-dimensional root finding so that the difference between the 97.5th and 2.5th percentiles equaled the target full width:

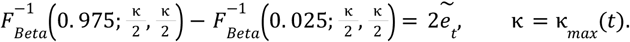

Smaller reference half-widths therefore produced larger reference concentrations. During training, κ_*max*_(*t*) served as a soft tier-specific threshold: concentrations above this value incurred a normalized squared penalty. The regularization weight was fixed at 0.1 across all experiments rather than tuned to individual benchmarks. The final objective was:

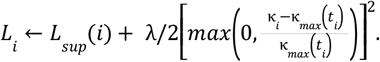

This source-aware regularization discourages implausibly narrow fitted distributions from noisy or weakly supervised data while preserving sharper distributions for high-quality quantitative measurements. Together, the beta likelihood, threshold-CDF supervision and source-aware concentration regularization enable PSICHIC+ to learn from large-scale heterogeneous bioactivity corpora without discarding noisy labels or collapsing all measurements into binary classes. The conditional beta distribution describes conditional response dispersion under point-estimated neural-network parameters; it does not capture epistemic uncertainty in model parameters, which would require ensembles, bootstrapping, or Bayesian inference.

### PSICHIC+ reparameterization of the beta distribution

The PSICHIC+ backbone encoder is identical to that of PSICHIC and computes the protein–ligand interaction fingerprint using physicochemical graph neural network layers. The difference lies in the prediction head. Whereas PSICHIC used task-specific MLPs to produce deterministic point outputs (a scalar p*K* for regression and logits for active/inactive classification), PSICHIC+ replaces these with two distributional MLP heads. From the interaction fingerprint, one head predicts an interior beta location μ *∈* (0, 1), and the other predicts a positive excess-concentration parameter φ > 0. We then reparameterize to shape parameters by adding a unit offset to guarantee α, β > 1, which restricts the fitted family to beta densities with a single interior mode:

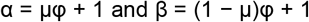

This yields a full conditional beta distribution, Beta(α, β), over normalized bioactivity. The total beta concentration is κ = α + β = φ + 2. The unit offset is a response-shape restriction, not a condition for numerical stability. Conceptually, μ defines the interior mode, while κ controls concentration: larger κ yields narrower conditional beta distributions around μ, whereas smaller κ yields broader distributions.

### Curating a 10^8^-scale bioactivity database

We assembled a large-scale, sequence-anchored bioactivity corpus based on Papyrus. Exact *K*_*i*_ and *K*_*d*_ measurements were retained as continuous pK labels, whereas *IC*_50_, *EC*_50_, and high-throughput screening records were converted to thresholded active or inactive labels. To provide negative supervision for each protein, we generated negative examples using three routes: (1) model-generated decoys,^43^ with five scaffold-diverse decoys sampled per active compound; (2) library decoys from ZINC20,^44^ filtered to remove close analogs of known actives; and (3) cross-target negatives, generated by pairing ligands with dissimilar proteins based on a Foldseek^45^ similarity threshold of less than 0.3. These generated and inferred negatives were assigned to separate source tiers and were not treated as experimental measurements.

Compound-target pairs were deduplicated and mapped to canonical SMILES and representative protein sequences. Each record was assigned to one of five source-quality tiers. Tier 1 comprised direct *K*_*i*_ and *K*_*d*_ measurements; Tier 2, thresholded labels derived from quantitative *IC*_50_ or *EC*_50_ measurements; Tier 3, high-throughput screening labels; Tier 4, model-generated decoys and cross-target negatives; and Tier 5, negatives sampled randomly from ZINC20. Tier 1 therefore contained continuous affinity measurements, whereas Tiers 2 through 5 contained binary labels with progressively weaker source reliability.

For each source tier *t*, we prespecified a reference half-width *h*_*t*_ for a central 95% predictive interval on the pK scale. The values of *h*_*t*_ were 1, 1.25, 1.5, 2 and 3 pK units for Tiers 1 through 5, respectively. We converted each value to the normalized scale as 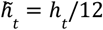 and defined κ_*max*_(*t*) using a symmetric beta reference distribution centered at 0.5. Specifically, κ_*max*_ (*t*) was obtained by one-dimensional root finding so that the difference between the 97.5th and 2.5th percentiles of *Beta*(κ/2, κ/2) equaled 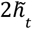 . Smaller reference half-widths therefore produced larger reference concentrations. During training, κ_*max*_ (*t*) served as a soft tier-specific threshold: concentrations above this value incurred a normalized squared penalty. The regularization weight λ was fixed at 0.1 in all experiments.

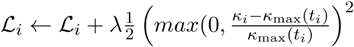

### Construction of the PSICHIC+ training sampling strategy

The PSICHIC+ training set was assembled from large-scale protein–ligand bioactivity data. Each training example consisted of a protein–small-molecule pair annotated with either a measured activity value or an inactive label. Measured affinity values were grouped by binding-strength class on a negative-log10 molar scale, using pK_I_/pK_D_, where larger values indicate stronger binding. Weak binders were defined as 4 ≤ pK_I_/pK_D_ < 6, corresponding approximately to >1–100 µM; moderate binders as 6 ≤ pK_I_/pK_D_ < 8, corresponding to >10 nM–1 µM; strong binders as 8 ≤ pK_I_/pK_D_ < 10, corresponding to >0.1–10 nM; and very strong binders as pK_I_/pK_D_ ≥ 10, corresponding to ≤0.1 nM. Active rows with potency values (IC_50_ or EC_50_) but without a quantitative affinity value were retained as qualitative binder examples. Inactive examples were retained as distinct evidence sources, including experimentally inactive measurements, high-throughput screening inactive calls, chemically generated decoys, random protein–ligand pairings and cross-target inactive pairs.

A naive sampling strategy that selected rows uniformly would overexpose proteins and assay sources with many database entries, particularly large inactive screens, while underexposing less-studied proteins and rarer active measurements (Supplementary Fig. 4). To reduce this bias, each training example was assigned a sampling weight that balanced exposure across three levels of the data: protein structural similarity, activity or inactivity type and Bemis–Murcko scaffold.

Proteins were first grouped by structural similarity using Foldseek clusters computed from AlphaFold DB v4^12^ predicted structures. We used the 90% Foldseek cluster assignment for the training table, so that proteins within the same structural cluster shared a single target-level exposure budget. For each structural cluster c, we calculated *N*_*c*_ as the number of distinct active scaffold–binding-strength observations represented in that cluster.

Each observation corresponded to a ligand scaffold paired with a binding-strength group. Thus, the same scaffold was counted separately if it was associated with multiple binding-strength groups; for example, a scaffold observed among both moderate and strong binders contributed two observations to *N*_*c*_ . The sampling budget for cluster c was set proportional to 1 + log(*N*_*c*_ ). This logarithmic scaling increased exposure for clusters with more active ligand evidence, while ensuring that additional observations produced progressively smaller increases in sampling weight. This retained information from well-studied target families while limiting over-representation of heavily assayed proteins.

Within each structural protein cluster, we used a deliberately active-enriched sampling scheme to emphasize experimentally informative binding and potency evidence while maintaining broad negative supervision. Active examples contain the graded signal needed to distinguish weak, moderate, strong and very strong binders, whereas inactive examples define the binder–non-binder boundary. For each cluster, 75% of the sampling budget was therefore assigned to active evidence and 25% to inactive evidence. The active budget was divided equally among the active evidence types present in the cluster, including qualitative binders and the measured affinity classes: weak, moderate, strong and very strong binders. The inactive budget was also stratified by source: high-throughput screening inactive examples received half of the inactive allocation, whereas ML-generated decoys, random protein–ligand pairing decoys and cross-target inactive pairing decoys shared the remaining half. This design preserved inactive examples as an important training signal while preventing any single negative source from dominating the batches.

Finally, within each activity or inactivity type, the assigned budget was balanced across chemical scaffolds. Each distinct scaffold within a given structural cluster and evidence type received an equal share of that type-specific budget. Rows with the same structural cluster, evidence type and scaffold then divided the scaffold-level budget equally. The final sampling probability of each training example was obtained by normalizing its row weight by the sum of all row weights. This procedure converted the raw database distribution into a balanced training-exposure distribution: the model continued to sample from the full corpus, but exposure was controlled across protein structural clusters, binding-strength classes, inactive evidence sources and molecular scaffolds.

### Zero-shot PSICHIC+–Boltz-2 ensemble with concentration-guided weighting

For each target, the PSICHIC+ and Boltz-2 predictions were independently oriented so that higher values indicated stronger predicted binding and were rescaled to the interval [0,1] across the candidate compounds.

For compound i, PSICHIC+ concentration was calculated as:

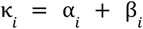

To reduce the influence of exceptionally large values, concentration was transformed using the natural logarithm and normalized within each target:

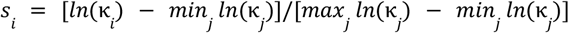

Here, j indexes the compounds evaluated for the target. We averaged an equal-weight prior, ω^*p*^ =0.5,, with a concentration-specific weight, 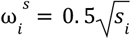 giving 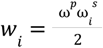. The final ensemble prediction was:

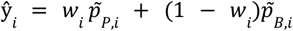

P and B denote the normalized PSICHIC+ and Boltz-2 predictions. PSICHIC+ provides both an activity prediction and a concentration signal that adapts its contribution to the ensemble for each compound. Predictions with higher concentration contribute more strongly, allowing the PSICHIC+ predictive distribution to guide the combined ranking without target-specific activity labels or retraining.

### PSICHIC+ few-shot recalibration of the conditional beta distribution

We use the pretrained PSICHIC+ predictive distribution as the starting point. A global calibration is estimated from the training subset, after which a local residual correction is applied most strongly when concentration is low and the ligand lies within the chemical space supported by the few-shot samples. Concretely, we first recalibrate the beta mean in logit space and temperature-scale the concentration:

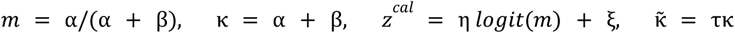

Here, η and ξ define the affine transformation of the logit mean, τ scales concentration, and all three hyperparameters are selected by grid search on the training subset. To capture local structure, we fit a local residual model Δ on the training set by regressing the logit residual *r=logit(y) −z* ^*cal*^ onto a ligand-similarity embedding, using weighted ridge regression. Training-example weights increase with pretrained concentration and local density, and fitted residuals are median-centered.

At test time, the fitted model predicts Δ, and we scale it using two multiplicative gates: a concentration gate 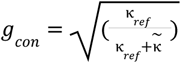 and a locality gate 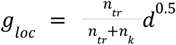 where *d* is the train-anchored similarity density computed from cosine similarities between ligand atom-pair count feature vectors, *n*_*tr*_ is the number of training compounds and *n* is a small constant. The final logit is:

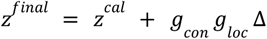

and the calibrated mean is µ^*final*^= σ(*z*^*final*^) . We propagate concentration by retaining 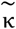. Unlike the pretrained prediction head, this post-hoc mean–concentration reconstruction is not restricted to α, β > 1. The final predictive beta parameters are:

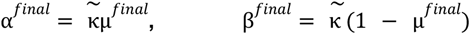

### PSICHIC+ traversal of chemical space via active learning

Whenever labels can be obtained, either from an available assay or from FEP calculations, PSICHIC+ ranks compounds using predicted activity and beta standard deviation. We support either a single-round selection (few-shot) or a multi-round active-learning loop when repeated propose-label-retrain cycles are feasible. In both settings, candidates are ranked by an acquisition score derived from the model’s predictive mean and beta standard deviation, and then down-selected to a chemically diverse batch.

PSICHIC+ produces, for each protein–ligand pair, beta distribution parameters (α, β) over a bounded activity target. From these, we compute the predictive mean (*U*) and beta standard deviation (σ) used to guide selection.

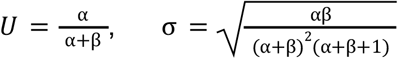

Acquisition scoring follows a Upper Confidence Bound (UCB)-style mean-plus-spread formulation with exploration coefficient λ. We use two regimes, greedy selection (λ = 0) and UCB selection (λ = 1):

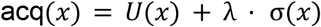

After ranking the unlabeled pool by acq(x), we restrict attention to the top 2 × *B* candidates, where *B* is the intended batch size. We then compute 2,048-bit ECFP4 fingerprints (radius 2), calculate pairwise Tanimoto distances and perform average-linkage hierarchical clustering into B clusters. The highest-scoring compound from each cluster is selected, yielding a diverse batch of B compounds. In the single-round selection mode, this batch is submitted for labeling and the process stops; in the multi-round (active-learning loop), newly acquired labels are appended, the model is updated by retraining, and the same propose-label-retrain cycle is repeated until the assay budget or timeline is exhausted.

### Benchmark evaluation and computational performance

Benchmark evaluations used the exclusion rules specified for each dataset. For DUD-E and DEKOIS 2.0, all PSICHIC variants were retrained separately after removing from pretraining any protein sharing at least 90% sequence identity with a benchmark target. Performance was calculated per target using AUROC, BEDROC (α = 80.5) and EF5%, and summarized across 102 DUD-E targets and 81 DEKOIS 2.0 targets. For the JACS and Merck FEP benchmarks, all benchmark ligands and proteins were excluded from pretraining. Pearson correlations were calculated within each of 16 target assays and summarized as the mean with s.e.m. Few-shot support sizes ranged from N = 1 to 6. Inference throughput was benchmarked on a single NVIDIA A100 GPU.

### PSICHIC+ optimization speedup

Inference throughput was benchmarked on a single NVIDIA A100 GPU using the FEP protein–ligand benchmark repeated ten times with randomized row order, yielding 4,610 input pairs. Throughput was reported as executed protein–ligand pairs per second. All reported optimization steps preserved the input row count, row order and row identifiers. For production-path measurements, every input row was passed through data loading and model forward execution; no exact-pair deduplication, output memoization or restored duplicate predictions were used. Warm-screening measurements excluded one-time graph construction where indicated, but retained molecular normalization, data loading, host-to-device transfer, model execution and output assembly unless otherwise specified.

#### PSICHIC and PSICHIC+ baseline

The first reference point was the published PSICHIC inference runtime of 0.008 s per protein–ligand pair, corresponding to 125 pairs per second. We used this value as the historical PSICHIC baseline. We then replayed the PSICHIC+ baseline on the A100 benchmark using a coding framework similar to that of PSICHIC. This baseline was slower than the PSICHIC reference because PSICHIC+ uses a more stringent molecular standardization path and produces additional distributional outputs, including beta-distribution parameters, beyond the original affinity prediction. These additions improve input consistency and provide distributional outputs but increase preprocessing, tensor handling and result-assembly cost before optimization.

Optimization was then performed as a sequence of modular NVIDIA GPU inference-system changes:

- <u>NVIDIA warm-cache inference:</u> First, we moved the benchmark into a controlled NVIDIA GPU container and used warm ligand and protein graph caches so that the timed run measured screening rather than repeated cold graph construction. This established a reproducible warm-cache PSICHIC+ inference harness.
- <u>Nsight-guided inference optimization:</u> Second, NVIDIA Nsight Systems profiling was used to identify host-side bottlenecks during inference. The dominant overheads were prediction extraction and row-order-preserving result assembly. We replaced row-wise output handling with a batched host-side inference fast path that preserved all protected outputs and the original input ordering.
- <u>CUDA inference graph optimization:</u> Third, we removed training-only graph operations from the inference path while keeping the pretrained PSICHIC+ weights fixed. In particular, MinCut loss and adjacency construction were bypassed during inference, and MotifPool was executed through its dense inference path. This reduced graph-execution overhead without changing the model parameters.
- <u>BioNeMo protein input reuse:</u> Fourth, we applied BioNeMo protein input reuse for screening workloads in which the same protein context is evaluated against multiple ligands. Protein-side dense input state was reused across rows with identical protein sequence context rather than reconstructed independently for each protein–ligand pair.
- <u>BioNeMo GPU protein cache:</u> Fifth, we extended protein reuse to a BioNeMo GPU protein graph-state cache. Protein graphs were first cached on disk. During inference, each row retained its original position and still entered the DataLoader and model forward path, but repeated protein tensors were represented by protein keys. Immediately before model execution, the required protein graph state was materialized from a bounded GPU cache. This avoided repeated CPU collation and repeated host-to-device transfer of identical protein graph tensors while preserving all-row execution.
- <u>NVIDIA Ampere TF32 mode:</u> Sixth, we evaluated NVIDIA Ampere TF32 execution. TF32 improved A100 throughput by using Ampere tensor-core arithmetic for compatible operations. Because TF32 introduced protected-output drift beyond the strict fp32 gate for alpha and beta distributional outputs, fp32 was retained as the strict reproducibility control, whereas TF32 was reported as the A100 throughput mode.

Across ten repeats of the FEP benchmark, throughput was 125 pairs per second for the historical PSICHIC baseline and 72.9 pairs per second for unoptimized PSICHIC+. The successive optimization stages increased PSICHIC+ throughput to 211.2, 237.7, 265.8, 276.9, 635.7 and 748.1 pairs per second.

## Data availability

The public datasets used in this study are available from the sources cited in the manuscript. The processed training and evaluation data needed to reproduce the analyses will be made available upon publication.

## Competing interests

J.B. and Y.-J.C. are employees of NVIDIA Corporation. All authors declare no other competing interests.

## Code availability

The source code for PSICHIC+ is available on GitHub at https://github.com/huankoh/psichic_plus

## Author contributions

H.Y.K. designed and developed the PSICHIC+ method. H.Y.K., H.J.W.K. and Y.Z. evaluated PSICHIC+ against leading methods for hit identification and hit-to-lead optimization. H.Y.K., H.J.W.K., Y.Z. and V.N. worked on the lab-in-the-loop framework and experiment. H.Y.K., J.B. and Y.-J.C. worked on optimizing the PSICHIC+ model and inference workflow using NVIDIA hardware and software tooling. D.S. contributed to the statistical framing, interpretation and revision of the manuscript. H.Y.K., A.T.N.N., S.P., L.T.M. and G.I.W. supervised the project. Project design, data interpretation and manuscript preparation were performed by all authors.

## Acknowledgements

This work was conducted while H.Y.K. was a PhD candidate supported by an Australian Government Research Training Program (RTP) Scholarship. The work was also supported by the Australian Research Council under grant DP210100072 and the National Health and Medical Research Council of Australia through Ideas Grant APP2013629. Some graphical elements in the figures were created with BioRender.com. Computational resources were provided by the Nectar Research Cloud, supported by the Australian Research Data Commons through the National Collaborative Research Infrastructure Strategy, and by the MASSIVE high-performance computing facility.

## References

1. Kitchen, D. B., Decornez, H., Furr, J. R. & Bajorath, J. Docking and scoring in virtual screening for drug discovery: methods and applications. Nat. Rev. Drug Discov. 3, 935–949 (2004).

2. Hopkins, A. L. Predicting promiscuity. Nature 462, 167–168 (2009).

3. Passaro, S. et al. Boltz-2: Towards Accurate and Efficient Binding Affinity Prediction. Preprint at 10.1101/2025.06.14.659707 (2025).

4. Lu, C. et al. OPLS4: Improving Force Field Accuracy on Challenging Regimes of Chemical Space. J. Chem. Theory Comput. 17, 4291–4300 (2021).

5. St. John, P. et al. BioNeMo Framework: a modular, high-performance library for AI model development in drug discovery. Preprint at 10.48550/arXiv.2411.10548 (2024).

6. Brenner, S. & Lerner, R. A. Encoded combinatorial chemistry. Proc. Natl. Acad. Sci. U. S. A. 89, 5381–5383 (1992).

7. Koh, H. Y., Nguyen, A. T. N., Pan, S., May, L. T. & Webb, G. I. Physicochemical graph neural network for learning protein–ligand interaction fingerprints from sequence data. Nat. Mach. Intell. 6, 673–687 (2024).

8. Mysinger, M. M., Carchia, M., Irwin, J. J. & Shoichet, B. K. Directory of Useful Decoys, Enhanced (DUD-E): Better Ligands and Decoys for Better Benchmarking. J. Med. Chem. 55, 6582–6594 (2012).

9. Bauer, M. R., Ibrahim, T. M., Vogel, S. M. & Boeckler, F. M. Evaluation and Optimization of Virtual Screening Workflows with DEKOIS 2.0 – A Public Library of Challenging Docking Benchmark Sets. J. Chem. Inf. Model. 53, 1447–1462 (2013).

10. Shen, C. et al. Boosting Protein–Ligand Binding Pose Prediction and Virtual Screening Based on Residue–Atom Distance Likelihood Potential and Graph Transformer. J. Med. Chem. 65, 10691–10706 (2022).

11. Cao, D. et al. Generic protein–ligand interaction scoring by integrating physical prior knowledge and data augmentation modelling. Nat. Mach. Intell. 6, 688–700 (2024).

12. Varadi, M. et al. AlphaFold Protein Structure Database in 2024: providing structure coverage for over 214 million protein sequences. Nucleic Acids Res. 52, D368–D375 (2024).

13. Chakravarty, D. et al. AlphaFold predictions of fold-switched conformations are driven by structure memorization. Nat. Commun. 15, 7296 (2024).

14. Škrinjar, P., Eberhardt, J., Tauriello, G., Schwede, T. & Durairaj, J. Have protein-ligand cofolding methods moved beyond memorisation? Preprint at 10.1101/2025.02.03.636309 (2025).

15. Masters, M. R., Mahmoud, A. H. & Lill, M. A. Investigating whether deep learning models for co-folding learn the physics of protein-ligand interactions. Nat. Commun. 16, 8854 (2025).

16. Burley, S. K. et al. Updated resources for exploring experimentally-determined PDB structures and Computed Structure Models at the RCSB Protein Data Bank. Nucleic Acids Res. 53, D564–D574 (2025).

17. Durairaj, J. et al. PLINDER: The protein-ligand interactions dataset and evaluation resource. Preprint at 10.1101/2024.07.17.603955 (2024).

18. Su, M. et al. Comparative Assessment of Scoring Functions: The CASF-2016 Update. J. Chem. Inf. Model. 59, 895–913 (2019).

19. Béquignon, O. J. M. et al. Papyrus: a large-scale curated dataset aimed at bioactivity predictions. J. Cheminformatics 15, 3 (2023).

20. Zdrazil, B. et al. The ChEMBL Database in 2023: a drug discovery platform spanning multiple bioactivity data types and time periods. Nucleic Acids Res. 52, D1180–D1192 (2024).

21. Abramson, J. et al. Accurate structure prediction of biomolecular interactions with AlphaFold 3. Nature 630, 493–500 (2024).

22. Sutton, R. The Bitter Lesson. Incomplete Ideas http://www.incompleteideas.net/IncIdeas/BitterLesson.html (2019).

23. Landrum, G. A. & Riniker, S. Combining IC50 or Ki Values from Different Sources Is a Source of Significant Noise. J. Chem. Inf. Model. 64, 1560–1567 (2024).

24. Feng, B. et al. A bioactivity foundation model using pairwise meta-learning. Nat. Mach. Intell. 6, 962–974 (2024).

25. Lu, W. et al. TANKBind: Trigonometry-Aware Neural NetworKs for Drug-Protein Binding Structure Prediction. Adv. Neural Inf. Process. Syst. 35, 7236–7249 (2022).

26. Miller, E. B. et al. Enabling structure-based drug discovery utilizing predicted models. Cell 187, 521–525 (2024).

27. Schneider, G. Automating drug discovery. Nat. Rev. Drug Discov. 17, 97–113 (2018).

28. Wang, L. et al. Accurate and Reliable Prediction of Relative Ligand Binding Potency in Prospective Drug Discovery by Way of a Modern Free-Energy Calculation Protocol and Force Field. J. Am. Chem. Soc. 137, 2695–2703 (2015).

29. Schindler, C. E. M. et al. Large-Scale Assessment of Binding Free Energy Calculations in Active Drug Discovery Projects. J. Chem. Inf. Model. 60, 5457–5474 (2020).

30. Finn, C., Abbeel, P. & Levine, S. Model-agnostic meta-learning for fast adaptation of deep networks. in Proceedings of the 34th International Conference on Machine Learning - Volume 70 1126–1135 (JMLR.org, Sydney, NSW, Australia, 2017).

31. Snell, J., Swersky, K. & Zemel, R. Prototypical networks for few-shot learning. in Proceedings of the 31st International Conference on Neural Information Processing Systems 4080–4090 (Curran Associates Inc., Red Hook, NY, USA, 2017).

32. Frey, N. C. et al. Lab-in-the-loop therapeutic antibody design with deep learning. Preprint at 10.1101/2025.02.19.639050 (2025).

33. Gorantla, R., Kubincová, A., Suutari, B., Cossins, B. P. & Mey, A. S. J. S. Benchmarking Active Learning Protocols for Ligand-Binding Affinity Prediction. J. Chem. Inf. Model. 64, 1955–1965 (2024).

34. Boby, M. L. et al. Open science discovery of potent noncovalent SARS-CoV-2 main protease inhibitors. Science 382, eabo7201 (2023).

35. Shen, W., Tang, H., Li, J., Li, X. & Chen, S. Multimodal data fusion for supervised learning-based identification of USP7 inhibitors: a systematic comparison. J. Cheminformatics 15, 5 (2023).

36. Zhang, Z., Zhao, B., Xie, A., Bian, Y. & Zhou, S. Activity Cliff Prediction: Dataset and Benchmark. Preprint at 10.48550/arXiv.2302.07541 (2023).

37. Thompson, J. et al. Optimizing active learning for free energy calculations. Artif. Intell. Life Sci. 2, 100050 (2022).

38. Shuker, S. B., Hajduk, P. J., Meadows, R. P. & Fesik, S. W. Discovering high-affinity ligands for proteins: SAR by NMR. Science 274, 1531–1534 (1996).

39. Backus, K. M. et al. Proteome-wide covalent ligand discovery in native biological systems. Nature 534, 570–574 (2016).

40. Buchwald, P. A Receptor Model With Binding Affinity, Activation Efficacy, and Signal Amplification Parameters for Complex Fractional Response Versus Occupancy Data. Front. Pharmacol. 10, 605 (2019).

41. Black, J. W. & Leff, P. Operational models of pharmacological agonism. Proc. R. Soc. Lond. B Biol. Sci. 220, 141–162 (1983).

42. Kenakin, T., Watson, C., Muniz-Medina, V., Christopoulos, A. & Novick, S. A Simple Method for Quantifying Functional Selectivity and Agonist Bias. ACS Chem. Neurosci. 3, 193–203 (2012).

43. Imrie, F., Bradley, A. R. & Deane, C. M. Generating property-matched decoy molecules using deep learning. Bioinformatics 37, 2134–2141 (2021).

44. Irwin, J. J. et al. ZINC20—A Free Ultralarge-Scale Chemical Database for Ligand Discovery. J. Chem. Inf. Model. 60, 6065–6073 (2020).

45. van Kempen, M. et al. Fast and accurate protein structure search with Foldseek. Nat. Biotechnol. 42, 243–246 (2024).

