## Supplementary Information for "Bioactivity foundation model for hit identification and hit-to-lead optimization from sequence data"

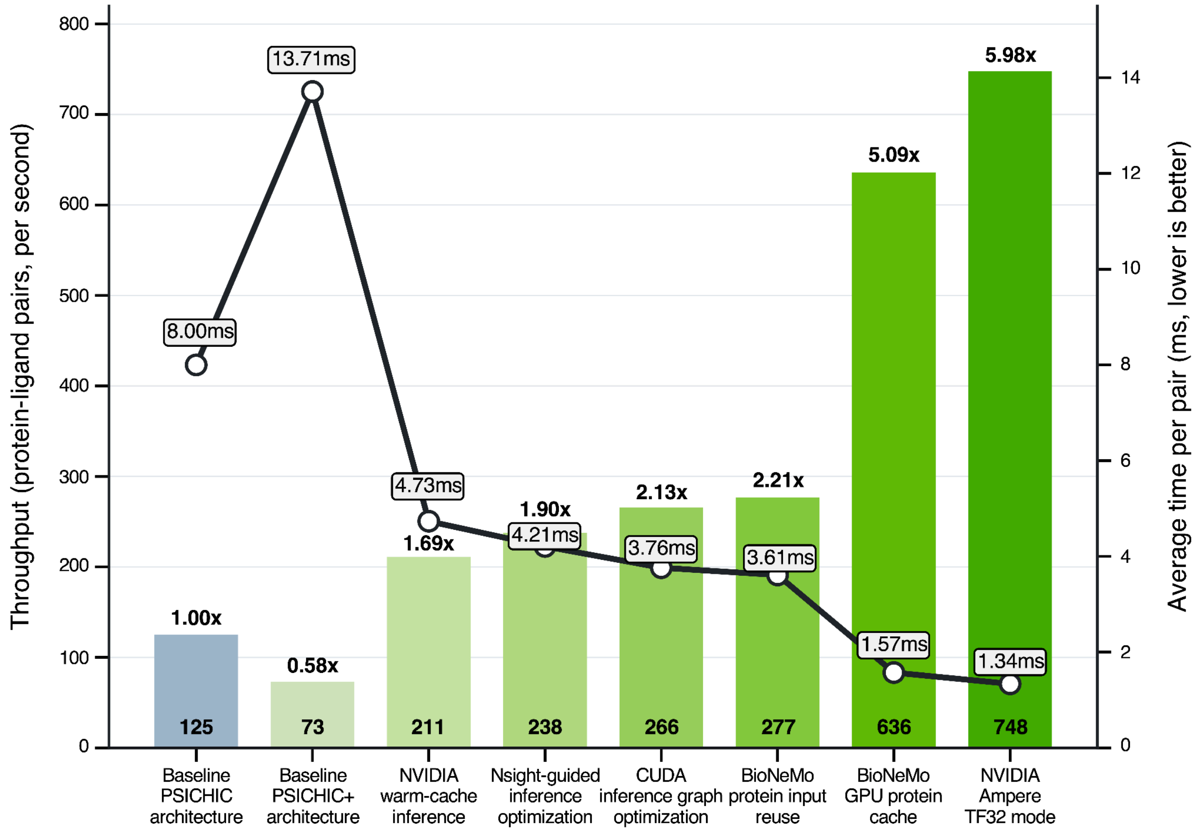


**Supplementary Fig. 1 | PSICHIC+ NVIDIA optimization speedup.** Stepwise optimization of PSICHIC+ inference throughput on a single NVIDIA A100 GPU. Bars indicate wall-clock screening throughput in protein–ligand pairs per second, and the black line indicates the corresponding average time per pair, with lower values indicating faster inference. The benchmark used the FEP+ dataset repeated ten times with randomized row order. The published PSICHIC runtime of 125 pairs/s, equivalent to 8.00 ms per pair, was used as the reference baseline. The unoptimized PSICHIC+ baseline measured 73 pairs/s, or 13.71 ms per pair, reflecting the additional cost of stricter molecular normalization and distributional output modeling, including beta-distribution parameters. Throughput was then improved through modular NVIDIA GPU inference-system optimizations: NVIDIA warm-cache inference, Nsight-guided inference I/O optimization, CUDA inference graph optimization, BioNeMo protein input reuse, BioNeMo GPU protein cache, and NVIDIA Ampere TF32 mode (see Methods). The final A100 TF32 configuration reached 748 pairs/s, or 1.34 ms per pair, corresponding to a 5.98-fold speedup over the PSICHIC baseline while preserving all-row execution without pair deduplication or output caching.

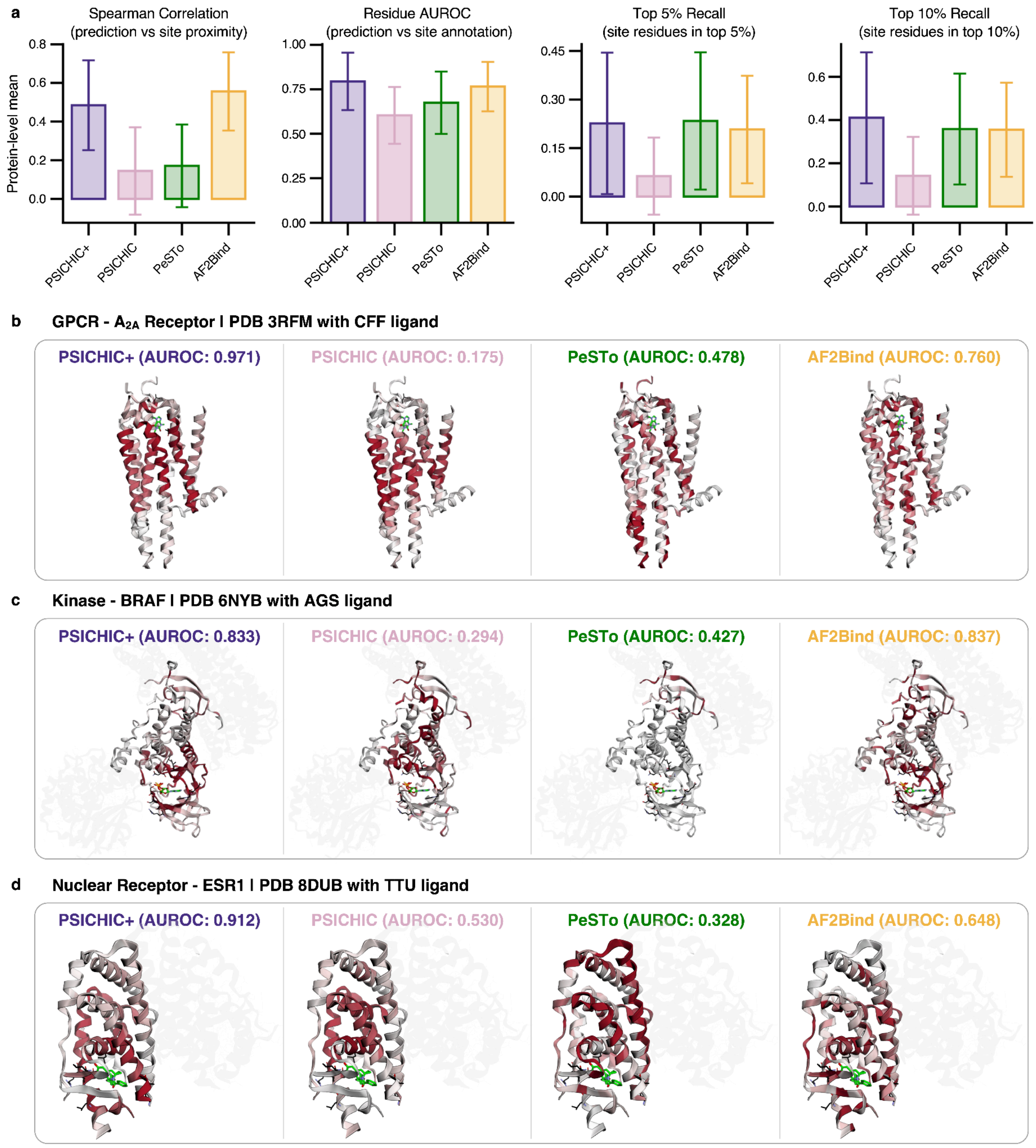


**Supplementary Fig. 2 | Comparison of residue-level binding-site predictions on BioLiP. a**, Performance of PSICHIC+, PSICHIC, PeSTo and AF2Bind. Spearman correlation measures the agreement between residue importance and proximity to the bound ligand. Residue AUROC measures the ranking of BioLiP binding-site residues relative to other residues. Top 5% and top 10% recall report the fraction of binding-site residues found among the highest-scoring residues. Bars show the mean across unique protein sequences and error bars show s.d. (n = 91,915). **b–d**, Residue-level predictions for $A_{2A}$ receptor bound to CFF (PDB 3RFM), BRAF bound to AGS (PDB 6NYB) and ESR1 bound to TTU (PDB 8DUB). Residue importance is colored from white (low) to dark red (high). Ligands are shown as green sticks, BioLiP binding-site residues as black sticks and other protein chains as translucent gray cartoons. AUROC values are reported for each complex. PSICHIC and PSICHIC+ use both the protein and ligand as input, whereas PeSTo and AF2Bind predict binding sites from the protein alone.

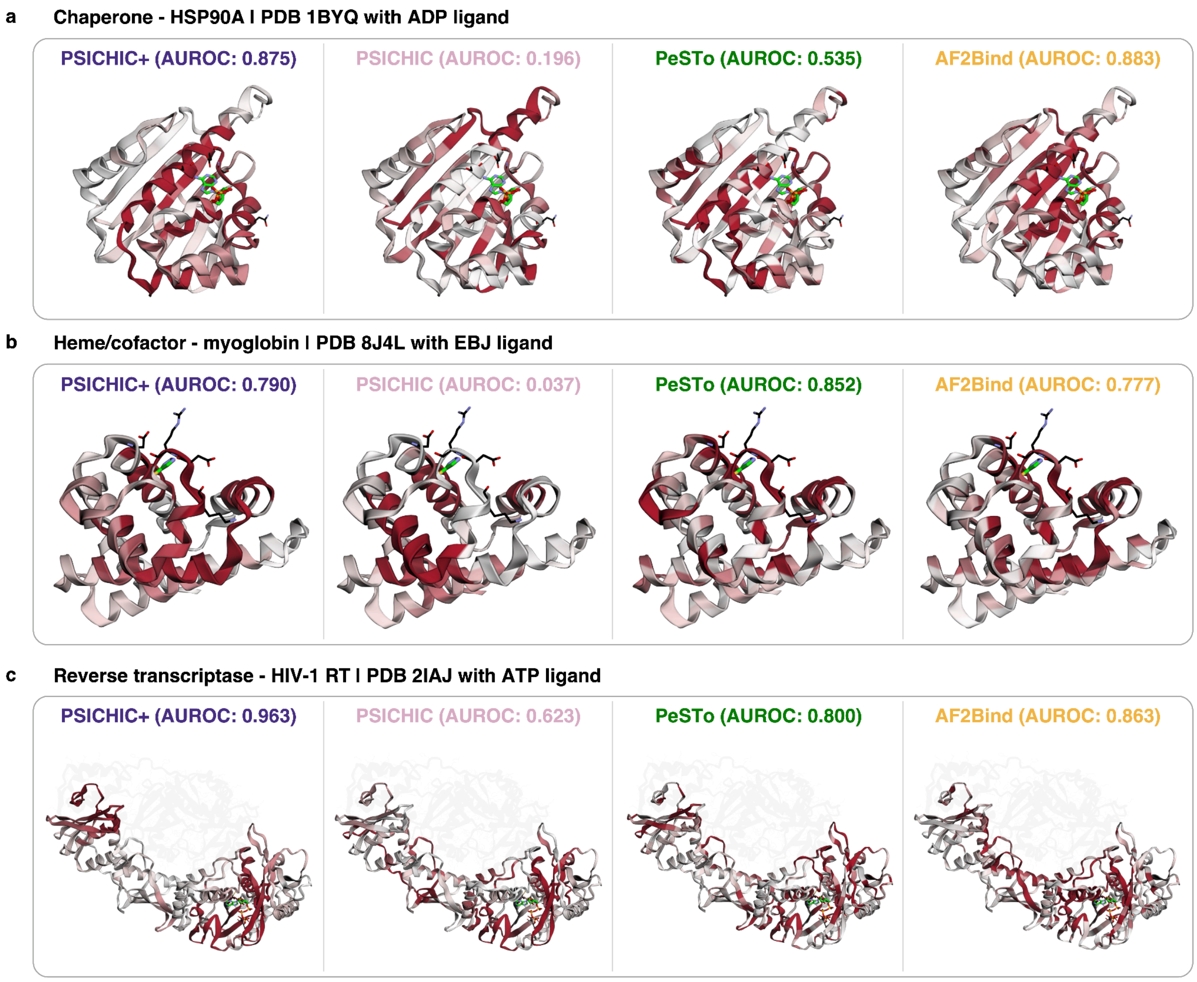


**Supplementary Fig. 3 | Residue-level predictions across additional protein families. a–c**, Comparison of PSICHIC+, PSICHIC, PeSTo and AF2Bind for HSP90A bound to ADP (PDB 1BYQ) (**a**), myoglobin bound to EBJ (PDB 8J4L) (**b**) and HIV-1 reverse transcriptase bound to ATP (PDB 2IAJ) (**c**). Residue importance is colored from white (low) to dark red (high). Ligands are shown as green sticks, BioLiP binding-site residues as black sticks and other protein chains as translucent gray cartoons. AUROC measures the ranking of annotated binding-site residues relative to other residues in the same protein. The same structure and orientation are used across methods for each complex.

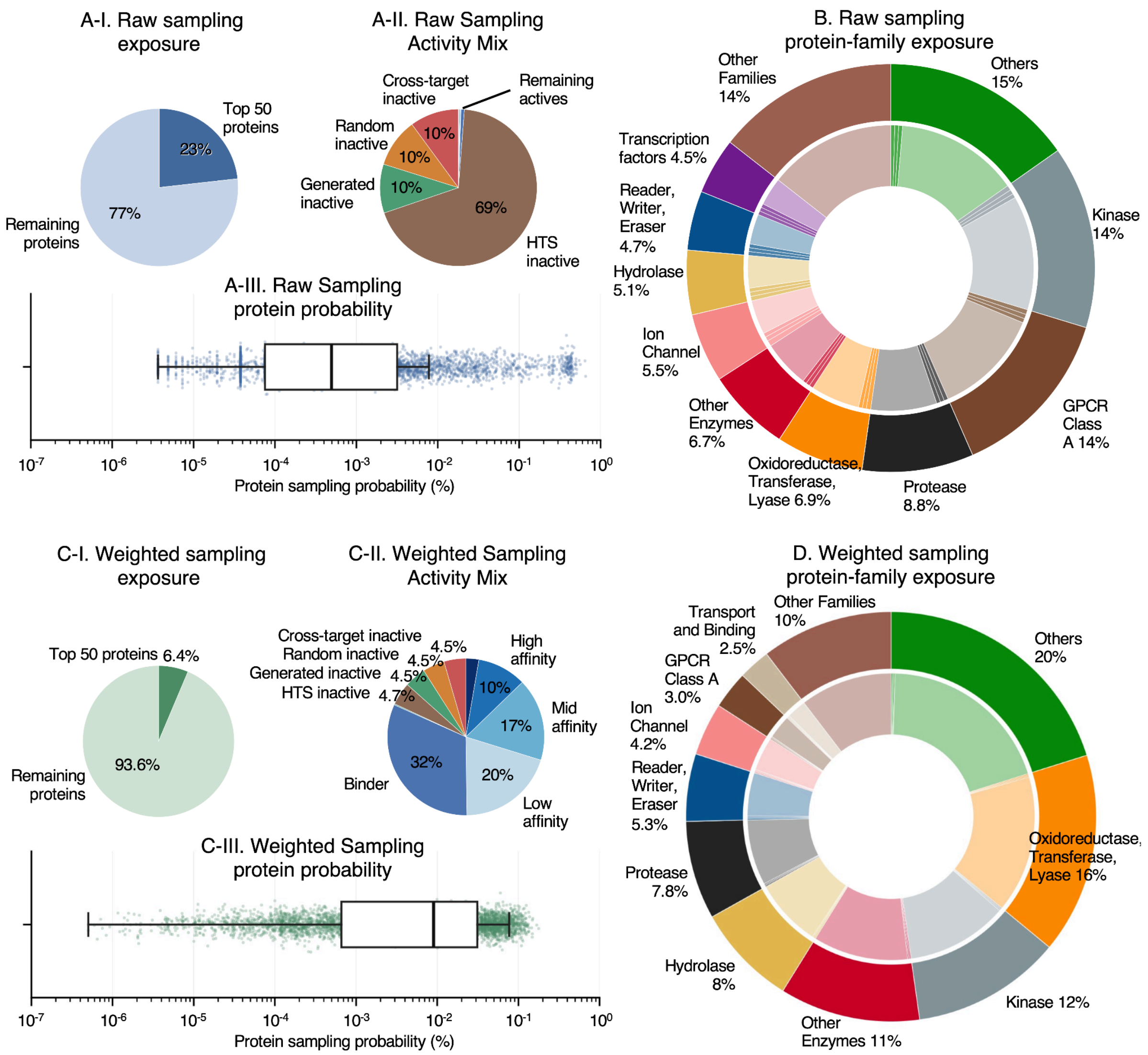


**Supplementary Fig. 4 | Broad protein-family and activity exposure in the PSICHIC+ training corpus.** The PSICHIC+ training corpus comprises approximately 82 million protein–ligand rows spanning 4,900 proteins and approximately 6.1 million unique ligands. **a**, Exposure under uniform row sampling, in which each protein–ligand row is sampled with equal probability. The panels show the contribution of the top 50 proteins, the activity/source composition and the resulting per-protein sampling probability, calculated as each protein's row count divided by the total number of training rows. **b**, Raw protein-family exposure under uniform row sampling. The outer ring shows family-level row fractions, and the inner ring shows the corresponding distribution across individual proteins within families. **c**, Exposure after applying the balanced training sampler described in Methods. Row weights reduce dominance by highly represented targets while preserving activity and source information. The panels show the weighted contribution of the top 50 proteins, the weighted activity/source composition and the resulting per-protein sampling probability, calculated as the sum of row weights for each protein divided by the total weight across all rows. **d**, Weighted protein-family exposure under the balanced sampler, shown as family-level exposure in the outer ring and individual-protein exposure in the inner ring. The sampler reduced the exposure of the top 50 proteins from 23.2% under uniform row sampling to 6.4% after weighting, while increasing effective exposure to qualitative binders and affinity-labeled active examples. This produced broader coverage across protein families, binding-strength regimes and inactive evidence sources during training.
